# Advanced Complexity and Plasticity of Neural Activity in Reciprocally Connected Human Cerebral Organoids

**DOI:** 10.1101/2021.02.16.431387

**Authors:** Tatsuya Osaki, Yoshiho Ikeuchi

**Affiliations:** Institute of Industrial Science, The University of Tokyo, Meguro, Tokyo 153-8505, Japan; Institute for AI and Beyond, The University of Tokyo, Bunkyo, Tokyo 113-8655, Japan

## Abstract

Macroscopic axonal connections in the human brain distribute information and neuronal activity across the brain. Although brain organoid technologies have recently provided novel avenues to investigate human brain function by replicating small neuronal networks of the brain *in vitro*, functionality of the macroscopic connections between distant regions has not been investigated with organoids. Here, we describe the neural activity of human cerebral organoids reciprocally connected by a bundle of axons. The connected organoids produced significantly more intense and complex oscillatory activity than conventional cerebral organoids. Optogenetic manipulations revealed that the connected organoids could respond to external stimuli and maintain the elevated frequency of neuronal activity for a short term, indicating that the connected organoids exhibit plasticity as a neuronal network. Our findings highlight functional importance of inter-regional connections and suggest that connected organoids can be used as a powerful tool for investigating the development and functions of macroscopic circuits in the human brain. can allow researchers to develop and test compounds to modulate activity of *in vitro* neuronal network models in unprecedented ways.

## Introduction

Organoid technologies have attracted increasing attention due to their unprecedented potential for modeling human organs *in vitro ^1–3^. In vivo*-like differentiation and self-organization can be achieved in organoids by mimicking *in vivo* development with growth factors and morphogens applied at appropriate timepoints and dosages. Organoids are expected to significantly promote neuroscience research, given the difficulties of investigating human brains noninvasively. Organoids that model brain regions including the cerebral cortex, thalamus, cerebellum, hippocampus, and choroid plexus have been reported ^4–10^ and offer new platforms to investigate these human brain regions *in vitro*. Brain organoids mimics structural and cellular characteristics of human brain development. They have been demonstrated to be competent research models for brain disorders including developmental and infectious disorders ^10–12^.

In addition to the structural and morphological similarities between organoids and the developing human brain, brain organoids have been used to model of human brain activity. Notably, recent studies have demonstrated that cerebral organoids exhibit robust oscillatory neuronal activity ^13^.

Cerebral organoids cultured for over 6 months begin to show oscillatory waves that resemble electroencephalography (EEG) activity patterns (delta band activity) of the preterm neonatal human brains. Further extension of the culture duration does not increase the complexity of neural activity ^13^, indicating that modeling the development of the brain with conventional organoids is intrinsically limited.

A key feature of the brain is the presence of macroscopic connections that bridge distant brain regions. Local circuitries in separated regions are interconnected by axons extended from one region to another, which enables a whole brain to function coordinately. The functionally and structurally distinct regions of the brain are often interconnected by reciprocal axonal projections that conduct action potentials ^14,15^, enabling information processing across multiple regions and subsequently higher cognitive function. Notably, individual organoids can only precisely model small regions of the brain; however, interactions between adjacent brain regions have been successfully modeled by fusing organoids that model two distinct brain regions ^16^. Therefore, models involving macroscopic axonal connections between distant regions, e.g., interconnected cortical areas positioned far apart, are required to advance brain organoid research. An ‘organoids-on-a-chip’ model of cerebral tracts has been generated by letting two cerebral organoids extend axons reciprocally in a ‘handshake’ manner to form a connection via a bundle of axons, similar to what occurs in the developing brain ^17^. In this model, a microfluidic compartment regulates the direction and assembly of growing axons, resulting in the formation of a thick axon bundle that connects two cerebral organoids. We hypothesized that these ‘connected organoids’ effectively model the axonal connections between physically distant brain regions by recapitulating the fundamental principles of macroscopic circuits and information processing. To test this hypothesis, we analyzed neuronal activity of the connected organoids generated from human induced pluripotent stem (iPS) cells on a multielectrode array and compared with single organoids and fused organoids. We observed highly complex activity in the connected organoids characterized by intense and irregular delta (0.5–4 Hz) and theta (4–8 Hz) components after culture for a relatively short-term duration (7–8 weeks). Optogenetic inhibition of neuronal activity of the axons between the organoids revealed that the connections between the organoids were critical for generating this complex activity. The activity of the connected organoids adapted to the temporal frequency of external stimulation, and activity was sustained at an elevated frequency after stimulation was terminated. Such adaptation of activity to temporal stimulation patterns required an induction delay for a few tens of seconds for the first stimulation event. Furthermore, the connected organoids regained the temporal patterns with a significantly reduced (less than ten seconds) when repeated stimulation was applied within a short time, indicating that the connected organoids exhibited short-term potentiation as an assembled macroscopic circuit. Thus, our findings highlight axonal connections between organoids significantly enhance functionality of cerebral organoids and support the importance of macroscopic connections within the brain, and the connected cerebral organoids as a promising tool to model macroscopic connections in functional neural circuits.

## Results and Discussion

### Reciprocally Connected Cerebral Organoids Cultured on a PDMS-MEA Chip Show Complex Neuronal Activity

To model the simplest type of macroscopic neural circuit, two regions connected by reciprocally projecting axons, we cultured a pair of human iPS cell-derived cerebral organoids in a microfluidic culture chip (**Fig. 1A and B**). The chip (PDMS-MEA chip) consisted of a multielectrode array (MEA) layer for recording neural activity, polydimethylsiloxane (PDMS) microfluidic layer for generating the connected organoids, medium reservoir ring for holding medium, and lid. The PDMS layer contained two holes, each of which received a cerebral organoid. The two holes were connected by a channel that provided spatial guidance for axons to reciprocally target the organoid at the opposite end of the channel. After 4 weeks of differentiation of cerebral organoids from human iPS cells, differentiation was confirmed by immunostaining and RT-PCR (**Fig. 1C and D**). Then they were introduced into the holes of the PDMS-MEA chip. Within 6 weeks, the two cerebral organoids were connected through an axon bundle in the chip (2 weeks in the chip, **Fig. 1E and Fig. S1**). The thickness of the axon bundle was approximately 75 μm after 6 weeks of differentiation and 120 μm after 8 weeks of differentiation (**Fig. 1F**). To assess ratio of excitatory and inhibitory neurons within the organoids, we performed immunostaining. vGlut1-positive excitatory neurons and GAD67-positive inhibitory neurons comprised approximately 70% and 5– 10% of cells in the organoids, respectively (**Fig. 1G**). The percentage of inhibitory neurons was lower than that of adult human brain (20-30%) ^18^, but comparable to mid-term human embryonic brains. Subcortical layers were observed in the connected organoids (**Fig. 1H**). These results indicate that the organoids mimicked developmental brain processes as they formed reciprocal axonal connections.

**Fig. 1.**
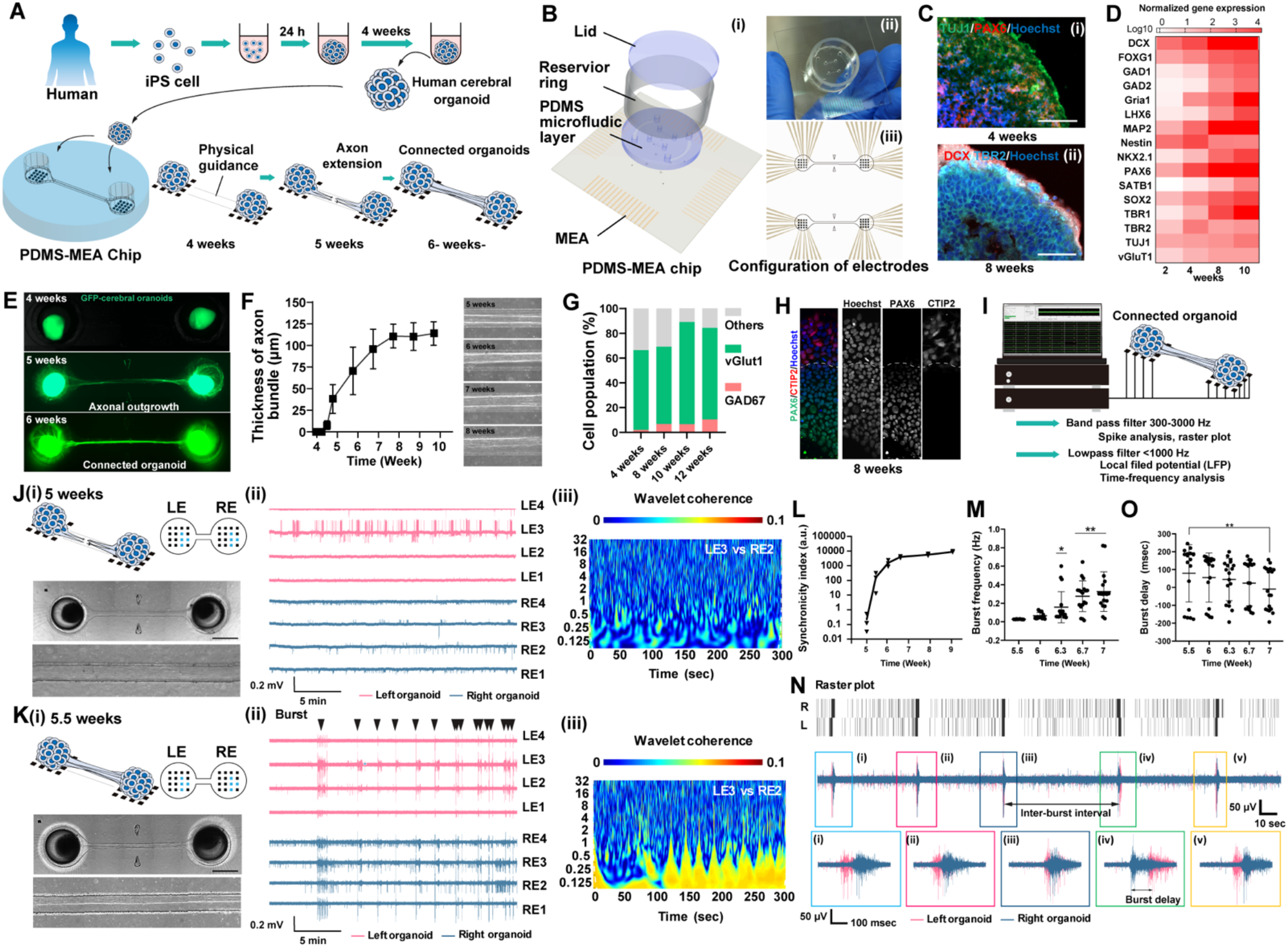
Formation and characterization of the connected organoids in a PDMS-MEA chip. (A) Schematic showing preparation of the connected cerebral organoids on a PDMS-MEA chip. One cerebral organoid was introduced into each of two chambers bridged by a microchannel in a chip. (B) The PDMS-MEA chip consisted of an MEA probe, patterned PDMS, a glass reservoir ring, and a PDMS lid (i, ii). Sixteen electrodes (in a four-by-four array) were located underneath each organoid (iii). (C) Representative internal structures after 4 and 8 weeks of culture. Scale bar: 150 μm. (D) Gene expression profiles in cerebral organoids from 2 to 10 weeks of culture. Normalized to GAPDH. (E) Axons projected from one organoid to another in 5 weeks, and thick axon bundles were formed by 6 weeks. (F) Axon bundle thickness over time. n = 6. (G) The proportions of excitatory neurons, inhibitory neurons and other neurons in the organoids. n = 3. (H) Immunohistochemical analyses revealed layers of different cell types within the connected organoids after 8 weeks of culture. Immunoreactivity against PAX6 and CTIP2 indicates the proliferative layer and cortical sublayer, respectively. (I) Recording of neuronal activity from the connected organoids in the chip. Raw analog signals from the electrodes were amplified and converted to digital signals (16 bit) at a 20-kHz sampling rate. Then, the signal was processed with a 300–3,000-Hz bandpass filter or a 1,000-Hz low-pass filter for spike analysis or local field potential (LFP) analysis, respectively. (J) Representative images of connected organoids after 5 weeks of culture. Scale bar: 1 mm (i). Filtered signals from four representative electrodes under each organoid (ii). Wavelet coherence between signals from an electrode under one organoid and another electrode under the other connected organoid (iii). (K) Representative images of the connected organoids after 5.5 weeks of culture. Scale bar: 1 mm (i). Synchronized burst-like activity associated with dense spikes was observed from multiple electrodes (ii). Wavelet coherence indicated a strong correlation between the two connected organoids (iii). (N) Synchronicity of activity in the two connected organoids increased during the culture period. (M) Burst frequency increased significantly with culture time. (N) Magnified view of the plot of neuronal activity of the two connected organoids. Synchronized burst activity was observed with a delay. (O) Burst delay after different culture periods. n = 20. *p<0.05, **p<0.01; one-way ANOVA. Error bars indicate the SD.

Electrodes within the MEA layer positioned under the two organoids captured neuronal activity (**Fig. 1B, I**). Action potential spikes and local field potentials (LFPs) were extracted by a high-frequency filter and a low-frequency filter, respectively, and assessed the time-course development. After 4.5-5 weeks of culture since starting differentiation (0.5-1 week of culture in the chip), electrical field potential was detected from the two cerebral organoids (**Fig. 1J**). At this stage, the activity of the two cerebral organoids was not synchronized, which was consistent with the lack of axonal connections between the organoids at this timepoint. At 5.5–6 weeks (1.5–2 weeks of culture in the chip), synchronized burst-like activity was observed (**Fig. 1K**). The upsurge and synchronization of this activity coincided with the physical connectivity of the organoids via an axon bundle. From 5 to 7 weeks of culture, neural activity became more synchronized, and more frequent burst-like activity was observed (**Fig. 1L and M**). Notably, a small temporal shift (around or less than 100 ms) between signals from the connected organoids was observed (**Fig. 1O and N**), suggesting that spontaneous neural activity was initiated in one organoid and propagated to the other organoid through the axon bundle. The two organoids alternated in their ability to initiate signal propagation, indicating that the connections were functionally bidirectional.

### Complex Neuronal Activity was Induced by Axonal Connections Between Cerebral Organoids

Next, we characterized developmental time-course of neuronal activity of the connected organoids. At 6 weeks (2 weeks of culture in the chip), slow LFP signals that had been absent at 4.5-5 weeks (prior to the establishment of axonal connections) were observed in the connected organoids (**Fig. 2A**). Clustered action potentials of multiple neurons are known to produce low-frequency LFP patterns in a coordinated manner ^19^. Consistently, after another week of culture in the chip (7 weeks of culture in total), the LFP patterns of the connected organoids became more complex, and intense activity in the delta frequency band (0.5–4 Hz) emerged (**Fig. 2A and B**), which indicates that coordinated ensemble of neuronal activity was developed. The intense and complex activity of the connected organoids at this relatively early stage was unexpected considering a previous report in which cerebral organoids exhibited delta band activity after culture for a few months ^13^.

**Fig. 2.**
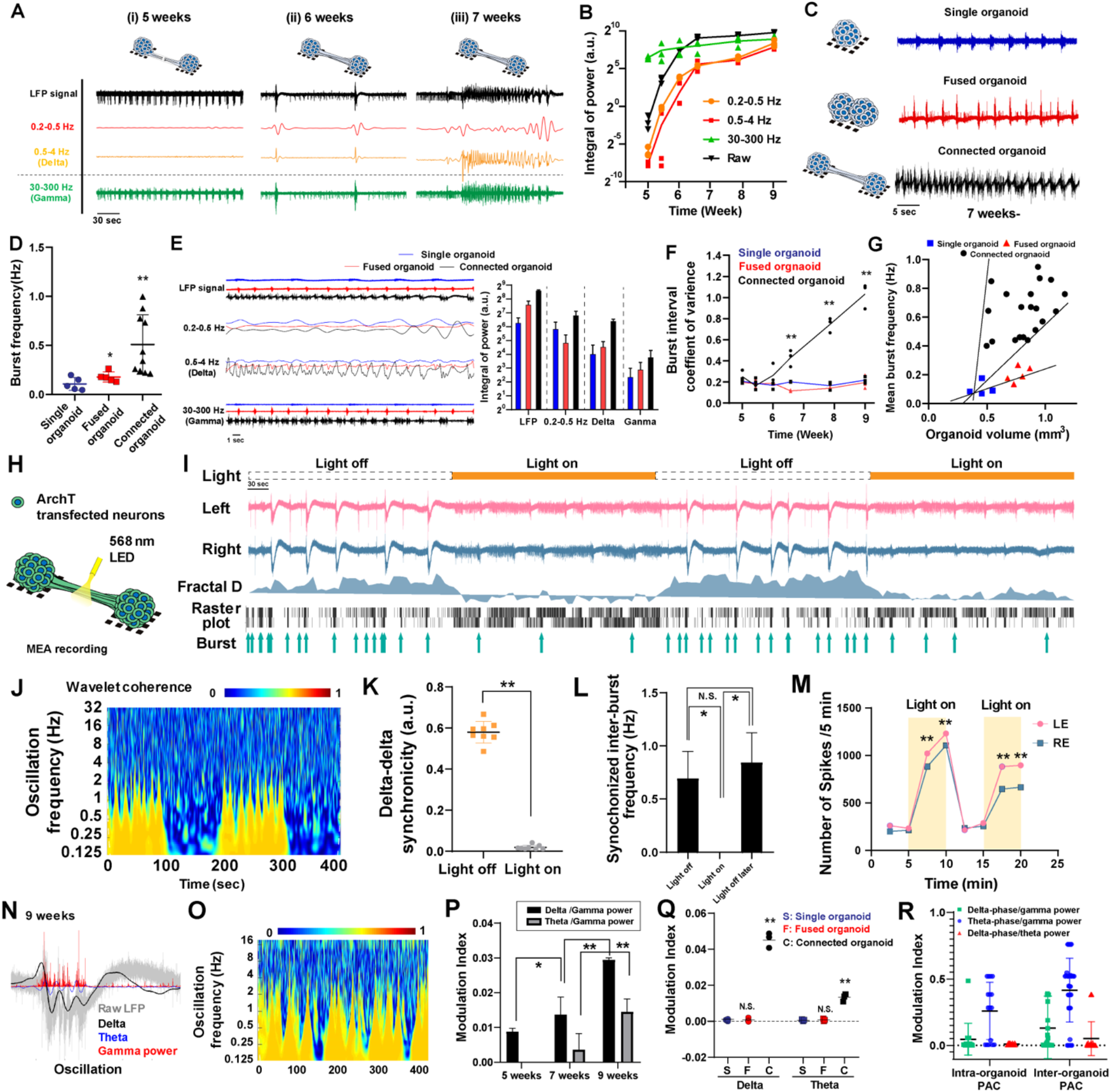
Reciprocal connections through bundled axons generate complex neuronal activity. (A) LFP signals were extracted from the 0.2–0.5-Hz, 0.5–4-Hz (delta), and 30–300-Hz (gamma) bands by inverse continuous wavelet transformation. At 8 weeks, the connected organoids generated slow-wave oscillations in the 0.5–4-Hz (delta) band. (B) Integral of power in wavelength bands. (C) Representative neuronal activity of the connected, single, and fused organoids. (D) Burst frequency of the connected, single, and fused organoids. n = 10. (E) Inverse continuous wavelet transformation in the 0.2–0.5-Hz, 0.5–4-Hz (delta), and 30–300-Hz (gamma) bands. Delta band oscillations were observed in the connected organoids but not in the single or fused organoids. (F) Interevent interval coefficient of variance among the three types of organoids. (G) Relationship between organoid volume and mean inter-burst frequency. (H) Setup for optogenetic inhibition of the synaptic interactions between two connected organoids via the inter-organoid axon bundle. (I) LFP and raster plot of the two connected organoids with and without light illumination. (J) Wavelet coherence revealed that slow-wave oscillations disappeared during light illumination, indicating the absence of inter-organoid correlated activity during light exposure. n = 8. (K) Inter-organoid synchronicity measured by delta phase-delta phase coupling was significant decreased during light illumination. (L) Light-induced inhibition of the axon bundle suppressed overall burst-like events. (M) The total number of action potential spikes over 5 min was quantified. Light exposure significantly increased isolated spikes. (N) Raw LFP plot from four electrodes of each of the connected organoids at 9 weeks. (O) Wavelet coherence between the two organoids showed synchronous activity in the theta band frequency. (P) Modulation index of phase-amplitude coupling in delta-phase/gamma-power and theta-phase/gamma-power of the connected organoids cultured for 5, 7, and 9 weeks. (Q) Delta-phase/gamma-power and theta-phase/gamma-power PAC modulation index of single, fused, and connected organoids. (R) Intra-or inter-organoid PAC modulation index in the connected organoids. *p<0.05, **p<0.01; one-way ANOVA. Error bars indicate the SD.

We thus compared the neuronal activity of our ‘connected’ organoids with that of conventional cerebral organoids (‘single’ organoids). Since the connected organoids contained approximately twice the number of cells, we also assessed the activity of tissues generated by directly fusing two cerebral organoids (‘fused’ organoids) as an additional control (**Fig. 2C and Fig. S2**). Two organoids were cultured together in a low-adhesion surface culture vessel to facilitate their spontaneous fusion. After 6 weeks of culture, oscillatory activity was detected in all the organoids (**Fig. 2C**). The burst-like activity was more frequent in the fused organoids than in the single organoids (**Fig. 2C**), suggesting that the total number of neurons in the organoids influenced neural activity. Notably, the connected organoids exhibited significantly more frequent burst-like activity than the single or fused organoids, although the numbers of neurons in the fused and connected organoids were comparable (**Fig. 2D**). These findings suggested that axonal connections between organoids strongly enhanced neural activity in the cerebral organoids.

Due to architectural significance of the axon bundle, we hypothesized that axon conduction delay mainly contributes to the induction of neuronal activity of the connected organoids. The signal propagation velocity between the fused organoids was significantly less than that within a single organoid, whereas the signal propagation velocity between the connected organoids was faster than that within a single organoid (**Fig. S2**). The lag in signal propagation between organoids was significantly higher in the fused organoids than in the connected organoids (**Fig. S2**). The axonal length did not alter the signal propagation velocity (**Fig. S3**), indicating that the synaptic connections within the organoids, instead of axon conduction delay, determined the signal propagation velocity. These results suggested that axonal connections served as a signaling ‘highway’ between the connected organoids and facilitated neuronal activity.

We then compared the neuronal activity of the single, fused, and connected organoids more detail in different frequency bands. The connected organoids exhibited significantly richer activity in the delta frequency band (0.5–4 Hz) than the single or fused organoids (**Fig. 2E**). To quantify the complexity of the temporal pattern of the neuronal activity in a different way, we assessed the periodicity of burst-like activity from the organoids. The coefficient of variance (CV) of burst-like events was significantly higher in the connected organoids than in the single or fused organoids after culturing more than 6 weeks (**Fig. 2F**), indicating that burst-like events were significantly less periodic and more intricate in the connected organoids than in the single or fused organoids. Specifically, the CV increased consistently over 9 weeks of culture in the connected organoids but did not change significantly in the single organoids or the fused organoids during this culture period. The fused organoids exhibited more frequent burst activity than the single organoids, consistent with their size (**Fig. 2G**), presumably due to the higher total numbers of cells and synapses within the assembled organoids. Strikingly, the connected organoids exhibited burst-like activity at a higher frequency than the single or fused organoids regardless of their size. These results suggested that the connected organoids recapitulated more complex neural circuits than the single or fused organoids and that the reciprocal connections induced complex activity in the cerebral organoids.

### Optogenetic Inhibition of Axon Bundles Between Connected Organoids

To test whether the axonal connections were critical for the complex activity of the connected organoids, we inhibited transmission through the axon bundle between the organoids with various approaches. Physically severing the axon bundle between organoids disrupted the burst-like events and LFP patterns in the delta band, highlighting the importance of the inter-organoid axonal connections for generating complex neuronal oscillatory activity patterns in the organoids (**Fig. S4**). To further assess the role of the axonal connections between the organoids, we modified the microfluidic chip to allow us to optogenetically inhibit the axon bundle between the organoids (**Fig. 2H and Fig. S5**). To achieve light-dependent inhibition of neuronal activity of the organoids, ArchT, an orange light-dependent outward proton pump ^20^, was expressed in organoids by infecting the organoids with an adeno-associated virus (AAV) vector before placing them in the PDMS-MEA chip. With an optic fiber and a PDMS lens, light was used to suppress neuronal activity in only the axon bundle in the microchannel. Illumination of the axon bundle with orange light (20 ms, 20 Hz for 5 min) robustly suppressed neuronal activity in the connected organoids, resulting in loss of high-amplitude burst-like events and low-frequency delta-LFP patterns under illumination (**Fig. 2I**). This finding suggested that action potentials traveling through the axon bundle induced burst-like activity in the connected organoids. Upon cessation of illumination, the burst-like activity and delta-LFP patterns were immediately restored. These light-induced responses were repeatedly observed after light exposure. The frequency of spontaneous burst-like events did not change after light exposure, suggesting that intrinsic internal circuit properties determined the frequency of the spontaneous burst-like activity.

Notably, the coherence and synchronicity of the signals from the two connected organoids were significantly degraded during light exposure (**Fig. 2J and K**). Indeed, optogenetic disconnection of the two organoids significantly suppressed the overall intensity of neural activity, potentially mimicking the effects of surgical treatments for refractory epilepsy patients 21,22. While burst-like events disappeared upon optogenetic inhibition of inter-organoid axons, the number of observed action potentials increased (**Fig. 2L and M**). This demonstrated that inter-organoid axons contributed to burst-like activity by temporally orchestrating and aggregating the activity of individual neurons within the two connected organoids. These results indicated that activity transmitted via the inter-organoid axon bundle underpinned complex neuronal activity in the connected organoids, consistent with the importance of macroscopic connections in the brain.

### Phase-Amplitude Coupling in the Connected Organoids

Extension of the culture period to 8 weeks resulted in a further increase in LFP frequency and action potential spikes of the connected organoids (**Fig. S6A**). The complexity of the signals also increased, and rich theta band activity was observed (**Fig. 2N**). The representative plot depicts a strong association of gamma activity with delta and theta activity. To examine the relationship between low-frequency activity and the amplitude of high-frequency spikes, phase-amplitude coupling (PAC), an established method to assess the relationship between complex EEG recordings in distinct frequency bands, was calculated ^23,24^. Within the burst-like oscillatory activity, LFP waves in the delta and theta bands coordinated with synchronized bursts of activity in the gamma band emerged (**Fig. 2N and O**). Delta-gamma PAC modulation of the connected organoids increased with culture time, followed by an increase in theta-gamma PAC modulation (**Fig. 2P**). Modulation index of both delta-gamma and theta-gamma in the connected organoids were significantly higher than those of the single or fused organoids (**Fig. 2Q**). Examination of each interconnected organoid in a pair revealed that delta-gamma PAC and theta-gamma PAC exhibited high modulation (intra-organoid PAC). The PAC between the organoids connected by the axon bundle (inter-organoid PAC) of both delta-gamma and theta-gamma modulations was higher than the intra-organoid PAC (**Fig. 2R**), indicating robust communication between the two organoids in the delta and theta bands. The correlated activity between the two organoids was validated using calcium imaging methods (**Fig. S7 and Movie S1**).

Next, to understand complexity of neuronal activity at the level of single spikes, we examined the neuronal avalanches of signals recorded from the connected organoids (**Fig. S8A**). Trains of temporally proximal signals that are no more than 2 ms apart across the electrodes were grouped and quantified as neuronal avalanches. Power law exponents of neuronal avalanches are considered a scale-free index of network critical dynamics ^25,26^. Avalanche size distributions increased over time as the connected organoids were cultured. At 5 weeks, the connected organoids exhibited an exponent of α = −2.8 (**Fig. S8B**). As the organoids grew axons in the channel and formed connections, the exponent increased to –2.1. At 8.5 weeks, the exponent became α = –1.6, which approximated the theoretical exponent of –3/2 in the critical branching process ^25,26^. We then examined the observed frequency of neuronal avalanches by pattern recognition using a hidden Markov model, which revealed that patterns of action potentials from the organoids in the chip increased over the culture time (**Fig. S8C**).

### Balanced Synaptic Receptors Underly the Complex Activity of the Connected Organoids

To investigate contribution of different types of synaptic receptors to the complex activity of the connected organoids, the connected organoids were treated with CNQX, APV, and bicuculline, antagonists of the major synaptic channels NMDA, AMPA, and GABA, respectively. No change in signal propagation speed was observed upon treatment with the antagonists (**Fig. S9**). However, the antagonists affected neuronal activity, as observed through action potential spikes and burst-like events (**Fig. S8D-G**). As expected, CNQX and APV decreased neuronal activity, delta-gamma PAC, and theta-gamma PAC in the connected organoids, suggesting that excitatory synaptic transmission played a critical role in driving complex activity in the connected organoids. Spiking activity was significantly increased, whereas the number of burst-like events remained unchanged following bicuculline treatment. However, the burst-like events became weaker following bicuculline treatment, suggesting that inhibitory synapses were critical for generating oscillatory activity in the connected organoids.

Next, we tested the effects of clinical compounds that function through modulating neuronal activities. Treatment with the GABA agonists baclofen and diazepam decreased the number of spikes, in accordance with the results obtained following treatment with bicuculline, which increased spiking activity. Treatment with an antipsychotic clozapine slightly decreased neuronal activity but did not alter PAC properties. Treatment with an opioid buprenorphine specifically decreased theta-gamma PAC, indicating that buprenorphine affected the coordination of activity in different frequency bands in the connected organoids. These results highlight the potential of using connected organoids to test the effects of compounds on complex activity.

### Short-term Facilitation Was Induced by Optogenetic Stimulation in the Connected Organoids

To characterize evoked responses of the connected organoids to external stimulations, we again employed an optogenetic approach. After expressing channelrhodopsin in the connected organoids, we stimulated the axons with pacing illumination (470 nm) at 0.5 Hz for 5 min, 1.0 Hz for 5 min, followed by 1.5 Hz for 5 min (**Fig. 3A**). Stimulation increased burst-like events in accordance with the temporal pattern of stimulation. After cessation of stimulation, the frequency of the burst-like events was sustained at a high level for more than 10 min but then returned to the prestimulation frequency (**Fig. 3B, C**). This sustained echo-like activity indicated that the temporal activity patterns of the connected organoids could be modulated by external stimulation and sustained for a certain period, highlighting the plasticity and capability of the connected organoids as a neuronal network. Notably, the induction of burst-like events occurred with a delay after the stimulation commenced (**Fig. 3C**), suggesting that multiple stimulation events were necessary to modulate the activity of the connected organoids. During this delay, the neuronal avalanches were extended in duration as the connected organoids were periodically stimulated (**Fig. 3D**), suggesting that the neurons in the connected organoids slowly adapted to the stimuli and established functional subcircuits before the connected organoids apparently followed the temporal patterns of external stimulation.

**Fig. 3.**
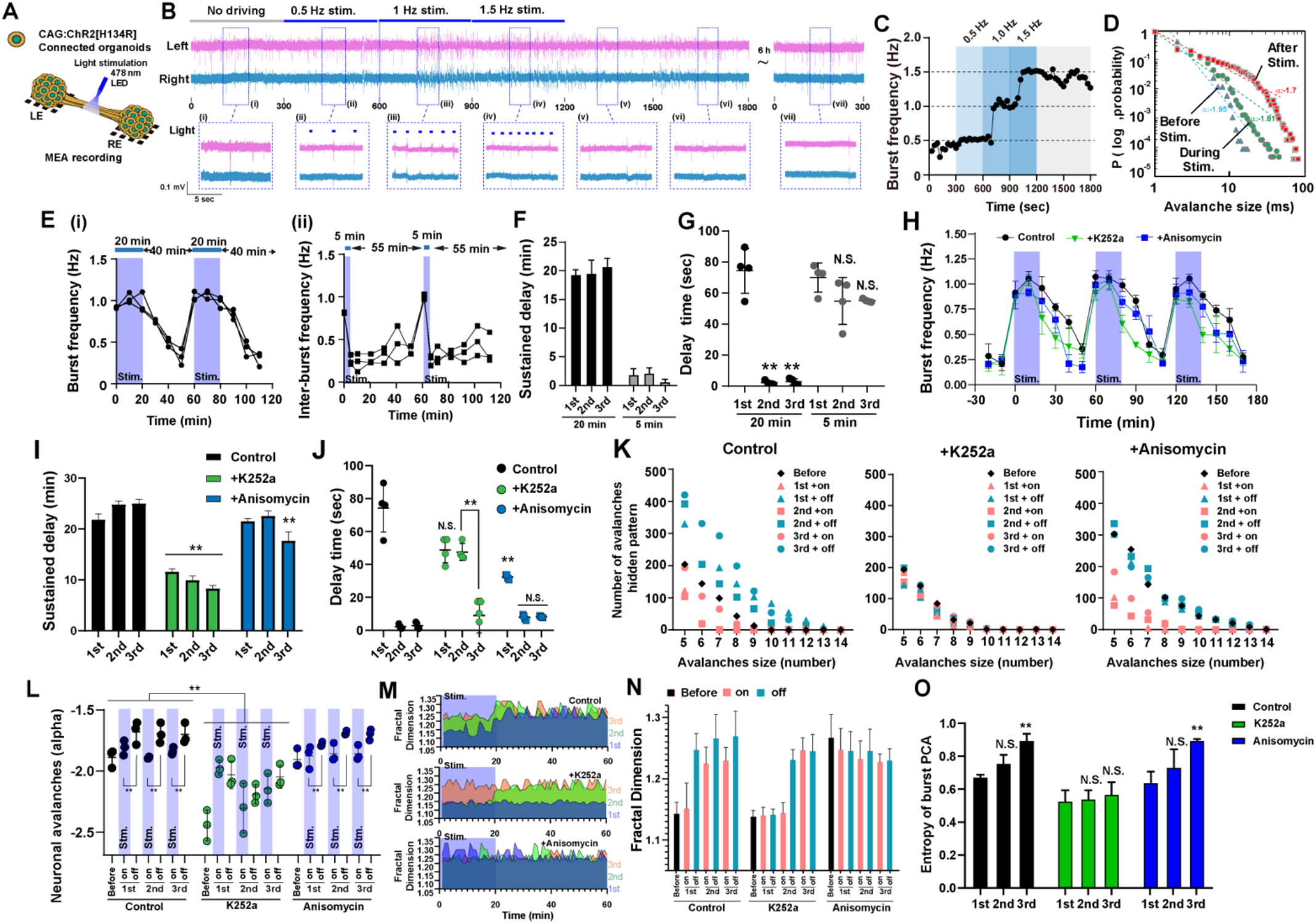
Potentiation of the connected organoids. (A) Optogenetic simulation of the axon bundle with a 470-nm laser drove synchronized burst activity at frequencies of 0.5, 1, and 1.5 Hz. The effect of stimulation persisted after cessation of illumination but did not persist 6 hours later. (B, C) Burst frequency was modulated by optical stimulation. The burst frequency followed the stimulation frequency after a significant delay. (D) Log plot of neuronal avalanche size and probability before, during, and after stimulation. (E) Time course of burst frequency with 1-Hz stimulation for 20 min (i) or 5 min (ii) every hour. (F) Duration until the burst frequency decreased to 75% of the maximum burst frequency after cessation of light stimulation. (G) The delay from the start of light stimulation to the induction of burst frequency was significantly reduced during the second and third attempts compared to that during the first attempt when the connected organoids were stimulated for 20 min. (H) Time series of burst frequency in the presence of K252a or anisomycin. (I) Duration of sustained activity of the connected organoids during treatment with the compounds. (J) Delay in the presence of K252a and anisomycin. (K) The number of hidden patterns in neuronal avalanches when treated with K252a or anisomycin. (L) Probability slope of neuronal avalanches. K252a treatment, but not anisomycin treatment, led to a decreased probability of neuronal avalanches. (M, N) FDs of the LFP signal. FDs increased after cessation of light stimulation during the first attempt under control conditions and during the second attempt upon K252a treatment. (O) Quantification of the diversity by calculating the entropy of burst PCA. n = 3. *p<0.05, **p<0.01; one-way ANOVA. Error bars indicate the SD.

Next, we examined the effects of repeated sessions of periodic optogenetic stimulation on the activity of the connected organoids. Repeated activation of neurons (‘rehearsal’) is known to maintain short-term memory in the human brain ^27^. Repeated stimulation resulted in echo-like activity after cessation of stimulation (20 min stimulation, **Fig. 3E and F**), while sustained activity was not observed with weak stimulation (5 min stimulation). Notably, the lag before the connected organoids adapted to the external stimulus was significantly shorter on the second and third attempts than on the first (**Fig. 3G**). This was observed only if the connected organoids were stimulated for 20 min, but not for 5 min. This indicated that the connected organoids were potentiated upon the first stimulation event and quickly regain the temporal pattern by adopting to the repeated stimulation.

Neuronal plasticity and memory are regulated by synaptic plasticity through diverse molecular programs, including calcium-dependent signaling pathways and local protein synthesis, essential for early and late phase of the response, respectively (refs). To probe the mechanisms underlying potentiation in the connected organoids, we treated the organoids with K252a, an inhibitor of the key calcium signaling protein CaM kinase II, and anisomycin, an inhibitor of protein synthesis, during the optogenetic stimulation experiment (**Fig. 3H**). The connected organoids responded to optical stimulation, and echo-like sustained activity after stimulation was observed upon treatment with K252a or anisomycin. The sustained period of poststimulation echo-like activity significantly decreased in the presence of K252a after the second and third stimulation events (**Fig. 3I**). In contrast, anisomycin treatment resulted in a slight decrease in echo-like activity after only the third stimulation event. After the second stimulation, K252a treatment inhibited the shortening of the lag between the start of stimulation and the response of the connected organoids compared to that of the control (**Fig. 3J**). These results indicated that calcium-dependent signaling pathways underpin the short-term potentiation observed in our experiments.

Neuronal avalanches also decreased in number with K252a treatment (**Fig. 3K**). During light exposure, the number of hidden patterns in neuronal avalanches decreased compared to that in the light-off period (**Fig. 3L and Fig. S10**). Notably, the number of hidden patterns gradually increased with repeated stimulation. Anisomycin treatment prevented the extension of hidden patterns for neuronal avalanches during the light-off period, suggesting that mechanisms of long-term potentiation were activated in the connected organoids.

Next, to examine the underlying variability and complexity of the activity of the connected organoids that are treated with K252a, anisomycin, or control, Higuchi’s fractal dimension (FD), which is utilized as an indicator of the complexity of EEG signals, was calculated ^28^. In the control, the FD increased after light stimulation ceased in the first stimulation event, and the elevated FD was preserved regardless of the second or the third light exposure (**Fig. 3M and N**). In the presence of K252a, FD did not increase in the first attempt but increased after light stimulation in the second attempt, and this increase was preserved in the third attempt, indicating that K252a treatment perturbed potentiation through complex network activity. In contrast, FD was not altered by light stimulation in the presence of anisomycin which elevated the FD baseline level with an unknown mechanism.

To assess temporal neuronal activity within burst-like activity, we sorted and aligned the evoked activity (**Fig. S10A**). Closer examination revealed that optogenetic stimulation events triggered multiple waves of neuronal activity within the evoked burst-like events. The latency of the burst-like events decreased with repeated rehearsal stimulation in the control (**Fig. S10B**). Such a decrease in activity was observed with anisomycin treatment but not in the presence of K252a. To dissect the evoked responses in further detail, we calculated the evoked burst probability histogram by kernel density estimation (**Fig. S10C and D**), which revealed that the first rise of the peak occurred earlier with repeated stimulation. Furthermore, the stimulation triggered secondary and tertiary waves of activity more than 200 ms after the primary response, and these waves of activity became more intense after repeated stimulation (**Fig. S10C and D**). Secondary waves were also observed with K252a or anisomycin treatment. Notably, two organoids interconnected via an axon bundle often responded to light exposure with slightly shifted kinetics (**Fig. S10E**). The secondary and tertiary waves of the evoked burst responses were found to alternate in the two connected organoids (**Fig. S10E**). These results suggested that the activity in the connected organoids was generated by a complex ensemble of activity in the two organoids. To quantify the divergence of the evoked activity, we evaluated the entropy of waveforms of the burst-like events from organoids treated with K252a or anisomycin and control organoids (**Fig. 3O**). Repeated stimulation increased the entropy of the evoked burst-like events; however, this effect was inhibited by K252a treatment, but not with anisomycin.

In conclusion, we demonstrate that connected cerebral organoids generate spontaneous high-frequency oscillations and are capable of exhibit plasticity. This indicate that inter-regional connections benefit functionality of neuronal circuits, and thus the importance of the inter-regional connections as fundamental architectural motif for the brain is recapitulated in the organoids-on-a-chip model. This system may facilitate the elucidation of human brain development and function from both physiological and pathological perspectives. To the best of our knowledge, the findings of this study provide the first evidence that cerebral organoids have capability to process and respond to external stimulation with variable short-term potentiation with repeated attempts. Currently, connected organoids can recapitulate only the primary response in signal processing. Further experiments such as connecting different regions of brain organoids and/or connecting more than two organoids via axon bundles will establish novel avenues for understanding the human brain. The results of such experiments would not only allow us to utilize this tissue model in biology but also provide insight into the development of novel algorithms in information processing.

## Materials and Methods

### Institutional Approval and Ethics

The use of human iPS cells was approved by Institute of Industrial Science, The University of Tokyo. The human iPS cells were handled in accordance with approved protocols.

### Experimental Design

#### Human iPS Cells

Human iPS cells were obtained from the Riken Cell Bank (409B2, HPS0076). The cells were maintained on ESC-qualified Matrigel-coated 6-well plates in mTeSR plus medium (STEMCELL Technologies) with 10 μM Y-23632 (only for the initial 24 hours; Wako) and subcultured every 5–7 days using ReLeSR reagent (STEMCELL Technologies).

#### Statistical Analysis

The reported values are the means of a minimum of three independent experiments. Data are presented as the mean ± SD. Comparisons were performed using one-way analysis of variance (ANOVA), with post hoc pairwise comparisons carried out using the Tukey-Kramer method. Statistical tests were performed using GraphPad Prism.

## Acknowledgments

We thank Drs. Yoji Hirano, Shoichiro Nakanishi, Yusuke Hirabayashi, Yoshito Masamizu, Parizad Billimoria, and Kazuyuki Aihara and the members of the Ikeuchi laboratory for discussion and critical comments. We thank Drs. Teruo Fujii, Soo Hyeon Kim, and Marie Shinohara for their generous support related to microfabrications.

## Author contributions

T.O. and Y.I. conceived of and designed the experiments. T.O. performed the experiments and analyzed the data. T.O. and Y.I. wrote the manuscript.

## Availability of data and materials

All data used to reach the conclusions in this paper are presented in the paper and Supplementary Materials. Additional data related to this paper may be requested from the authors.

## References

1 Kim, J., Koo, B. K. & Knoblich, J. A. Human organoids: model systems for human biology and medicine. Nat Rev Mol Cell Biol 21, 571–584, doi:10.1038/s41580-020-0259-3 (2020).

2 Dutta, D., Heo, I. & Clevers, H. Disease Modeling in Stem Cell-Derived 3D Organoid Systems. Trends Mol Med 23, 393–410, doi:10.1016/j.molmed.2017.02.007 (2017).

3 Rossi, G., Manfrin, A. & Lutolf, M. P. Progress and potential in organoid research. Nat Rev Genet 19, 671–687, doi:10.1038/s41576-018-0051-9 (2018).

4 Del Dosso, A., Urenda, J.-P., Nguyen, T. & Quadrato, G. Upgrading the Physiological Relevance of Human Brain Organoids. Neuron 107, 1014–1028, doi:https://doi.org/10.1016/j.neuron.2020.08.029 (2020).

5 Xiang, Y. et al. Fusion of Regionally Specified hPSC-Derived Organoids Models Human Brain Development and Interneuron Migration. Cell Stem Cell 21, 383–398.e387, doi:https://doi.org/10.1016/j.stem.2017.07.007 (2017).

6 Sakaguchi, H. et al. Generation of functional hippocampal neurons from self-organizing human embryonic stem cell-derived dorsomedial telencephalic tissue. Nature Communications 6, 8896, doi:10.1038/ncomms9896 (2015).

7 Muguruma, K., Nishiyama, A., Kawakami, H., Hashimoto, K. & Sasai, Y. Self-Organization of Polarized Cerebellar Tissue in 3D Culture of Human Pluripotent Stem Cells. Cell Reports 10, 537–550, doi:https://doi.org/10.1016/j.celrep.2014.12.051 (2015).

8 Xiang, Y. et al. hESC-Derived Thalamic Organoids Form Reciprocal Projections When Fused with Cortical Organoids. Cell Stem Cell 24, 487–497.e487, doi:https://doi.org/10.1016/j.stem.2018.12.015 (2019).

9 Pellegrini, L. et al. Human CNS barrier-forming organoids with cerebrospinal fluid production. Science 369, eaaz5626, doi:10.1126/science.aaz5626 (2020).

10 Lancaster, M. A. et al. Cerebral organoids model human brain development and microcephaly. Nature 501, 373–379, doi:10.1038/nature12517 (2013).

11 Song, E. et al. Neuroinvasion of SARS-CoV-2 in human and mouse brain. bioRxiv, doi:10.1101/2020.06.25.169946 (2020).

12 Krenn, V. et al. Organoid modeling of Zika and herpes simplex virus 1 infections reveals virus-specific responses leading to microcephaly. Cell Stem Cell, doi:10.1016/j.stem.2021.03.004 (2021).

13 Trujillo, C. A. et al. Complex Oscillatory Waves Emerging from Cortical Organoids Model Early Human Brain Network Development. Cell Stem Cell 25, 558–569.e557, doi:https://doi.org/10.1016/j.stem.2019.08.002 (2019).

14 Zingg, B. et al. Neural networks of the mouse neocortex. Cell 156, 1096–1111, doi:10.1016/j.cell.2014.02.023 (2014).

15 Swanson, L. W., Hahn, J. D. & Sporns, O. Organizing principles for the cerebral cortex network of commissural and association connections. Proc Natl Acad Sci U S A 114, E9692–E9701, doi:10.1073/pnas.1712928114 (2017).

16 Bagley, J. A., Reumann, D., Bian, S., Lévi-Strauss, J. & Knoblich, J. A. Fused cerebral organoids model interactions between brain regions. Nature Methods 14, 743–751, doi:10.1038/nmeth.4304 (2017).

17 Kirihara, T. et al. A Human Induced Pluripotent Stem Cell-Derived Tissue Model of a Cerebral Tract Connecting Two Cortical Regions. iScience 14, 301–311, doi:10.1016/j.isci.2019.03.012 (2019).

18 Markram, H. et al. Interneurons of the neocortical inhibitory system. Nat Rev Neurosci 5, 793–807, doi:10.1038/nrn1519 (2004).

19 Buzsáki, G., Anastassiou, C. A. & Koch, C. The origin of extracellular fields and currents--EEG, ECoG, LFP and spikes. Nat Rev Neurosci 13, 407–420, doi:10.1038/nrn3241 (2012).

20 Han, X. et al. A high-light sensitivity optical neural silencer: development and application to optogenetic control of non-human primate cortex. Front Syst Neurosci 5, 18, doi:10.3389/fnsys.2011.00018 (2011).

21 Spencer, S. S. et al. Corpus callosotomy for epilepsy. I. Seizure effects 38, 19–19, doi:10.1212/wnl.38.1.19 (1988).

22 Paul, L. K. et al. Agenesis of the corpus callosum: genetic, developmental and functional aspects of connectivity. Nature Reviews Neuroscience 8, 287–299, doi:10.1038/nrn2107 (2007).

23 Fell, J. & Axmacher, N. The role of phase synchronization in memory processes. Nat Rev Neurosci 12, 105–118, doi:10.1038/nrn2979 (2011).

24 Canolty, R. T. & Knight, R. T. The functional role of cross-frequency coupling. Trends Cogn Sci 14, 506–515, doi:10.1016/j.tics.2010.09.001 (2010).

25 Beggs, J. M. & Plenz, D. Neuronal Avalanches in Neocortical Circuits. The Journal of Neuroscience 23, 11167–11177, doi:10.1523/jneurosci.23-35-11167.2003 (2003).

26 Bowen, Z., Winkowski, D. E., Seshadri, S., Plenz, D. & Kanold, P. O. Neuronal Avalanches in Input and Associative Layers of Auditory Cortex. Frontiers in Systems Neuroscience 13, doi:10.3389/fnsys.2019.00045 (2019).

27 Himmer, L., Schönauer, M., Heib, D. P. J., Schabus, M. & Gais, S. Rehearsal initiates systems memory consolidation, sleep makes it last. Sci Adv 5, eaav1695, doi:10.1126/sciadv.aav1695 (2019).

28 Varley, T. F. et al. Fractal dimension of cortical functional connectivity networks & severity of disorders of consciousness. PLOS ONE 15, e0223812, doi:10.1371/journal.pone.0223812 (2020).

