## Supplemental Figures and Methods for "Advanced Complexity and Plasticity of Neural Activity in Reciprocally Connected Human Cerebral Organoids"

### Complex Activities and Short-term Memories in Reciprocally Connected Cerebral Organoids

**This PDF file includes:**

Materials and Methods
Figs. S1 to S10
Tables S1
Caption for Movie S1

**Other Supplementary Materials for this manuscript include the following:**

Movie S1

#### Materials and Methods

##### *PDMA-MEA chip fabrication*

SU-8 master positive patterned molds were fabricated by standard photolithography techniques described elsewhere (*20*). Briefly, SU-8 (2100 or 2075) was poured onto a silicon wafer (4 inches) and spin-coated (1200-1500 rpm, for 30 sec). The wafer was pre-baked for 9 min at 65°C and for 40 min at 95°C on a hotplate. Then, UV light (365 nm, 2.5-3.0 $mWcm^2$) was exposed with a photomask for 60-75 sec. The wafer was baked for 7 min at 65°C and for 13 min at 95°C on a hotplate. After cooling down the wafer, SU-8 was developed by using SU-8 developer for 15 min, then washed with isopropyl alcohol (IPA) for three times. The wafer was hard-baked for 3 min at 150°C in the oven. The thickness of the SU-8 was approximately 150 µm.

Microfluidic device was made with a polydimethylsiloxane (PDMS) silicone elastomer kit Sylgard184 (Dow Corning). Silicone elastomer and a curing agent were mixed at a weight ratio of 10:1, degassed, poured onto the patterned SU-8 structures, and cured in the oven at 80 °C for 6 h. The holes for organoids and reference electrode were created with biopsy punches (1.5 mm and 2 mm, respectably). A glass ring (inner diameter: 22 mm, outer diameter: 25 mm) as the medium reservoir was glued to the PDMS device. The devices were sterilized by autoclave, 70% ethanol, and UV treatment.

##### *Cerebral organoids formation*

To generated cerebral organoid, first, iPS cells were dissociated into single cells with TypLE express. Then, 20, 000 cells were plated to each well of U-bottom ultra-low attachment 96 well plate (Prime surface, Sumitomo bakelite) in mTeSR plus with 10 µM of Y-23632. After 24 h, culture medium was replaced with neural induction medium (DMEM-F12, 15% (v/v) knockout serum replacement, 1% (v/v) MEM-NEAA, 1% (v/v) Glutamax, 100 nM LDN-193189, and 10 mM SB431542) and the medium was changed every 2 days. After 10 days of culture, the culture medium was replaced with 1:1 mixture of DMEM/F12 and Neurobasal medium supplemented with 0.5% (v/v) N2 supplement, 1% (v/v) B27 supplement without vitamin A, 1% (v/v) Glutamax, 0.5% (v/v) MEM-NEAA, 0.25 mg/ml (v/v) human insulin solution, and 1% (v/v) Penicillin/Streptomycin and the medium was changed every 2 days until 18 days. After 18 days of culture, the culture medium was replaced with maintenance medium (Neurobasal medium supplemented with 0.5% (v/v) N2 supplement, 1% (v/v) B27 supplement with vitamin A, 1% (v/v) Glutamax, 0.5% (v/v) MEM-NEAA, 0.25 mg/ml (v/v) human insulin solution, 20 ng/ml BDNF, and 200 mM ascorbic acid, and 1% (v/v) Penicillin/Streptomycin). Cerebral organoids were cultured for four weeks and subjected to the connected organoid formation.

##### *Connected organoids formation in the microfluidic chip*

Two cerebral organoids were connected in the microfluidic device. The previously reported protocol (*13*) has been modified. Briefly, the PDMS devices were bonded to MEA probes. PDMS device should be positioned to align with the electrodes. The microchannel was coated with ESC-qualified Matrigel (Corning) in DMEM/F12 (1:30) for 1 hour at RT. The coating solution was replaced with maintenance medium. Cerebral organoids were then placed into the 1.5 mm of holes and settled down to the bottom by the gravity. The maintenance medium was replaced every 2 days in the microfluidic device.

##### *Multi-electrode array measurement and post-analysis*

In order to capture neuronal activity via multi-electrode array, maintenance medium was replaced with Brainphys supplemented with 1% (v/v) B27 supplement with vitamin A, 1% (v/v) Glutamax, 20 ng/ml BDNF, and 1%(v/v) Penicillin/Streptomycin before 24 h measurement. The PDMS-MEA chip was set to the MED64 system (Alpha MED Scientific) and electrical signals from all 64 electrodes were recorded for 5-30 min at 37°C at 20,000 Hz sampling rate. The recording noise was eliminated by band-pass filer between 0.1-10000 Hz during the measurement. The raw signal was further filtered by a bandpass filter (300-3000 Hz) for spike analysis, raster plot, and spike clustering, and low pass filter (<1000 Hz) for local field potential analysis, and then all post-analysis was performed using Signal Processing Toolbox, Curve fitting Toolbox, Deep learning Toolbox, Parallel Computing Toolbox, Wavelet Toolbox in MATLAB. All the analysis and calculation were conducted using MATLAB software. All the script for calculation in this paper was deployed and can be downloaded from https://github.com/TatsuyaOsaki/Matlab_function

##### *Wavelet coherence and Wavelet transformation for frequency isolation*

Wavelet coherence is a measure of correlation between two signals at specific frequency. Wavelet coherence from LFP recording, f(t) was calculated using function cwt() and icwt() in package "Wavelet Toolbox":

$$W(b,a) = \frac{1}{\sqrt{a}}\int_{-\infty}^{\infty} f(t)G\left(\frac{t-b}{a}\right)dt \quad (1)$$

where a, b denoted the scaling factor (1/Hz) and the center location (ms) of the mother wavelet function, respectively. In Equation (1), G(x) is the complex Morlet function:

$$G(x) = \frac{1}{\sqrt{\pi F_B}} exp\left(-\frac{x^2}{F_B}\right) exp(2i\pi F_c x) \quad (2)$$

where frequency bandwidth $F_B$ was set to 5, and the center frequency Fc was set to 1.

##### *Cross-correlation*

Cross-correlation $R$ was calculated with xcorr() function in MATLAB. Cross-correlation sequence of two signal series $x_n$ and $y_n$, is given by

$$R_{xy}(m) = E\{x_{n+m}y_n^*\} \; = \; E\{x_n y_{n-m}^*\}$$

where m is time lag, and $E$ is the expected value operator. We used scaleopt option to normalize the cross-correlation as 1 when two signals had no time lag:

$$\widehat{R}_{xycoeff}(m) = \frac{1}{\sqrt{\widehat{R}_{xx}}(0)\sqrt{\widehat{R}_{yy}}(0)})\sqrt{\widehat{R}_{xy}}(m)$$

##### *Neuronal avalanches*

Neuronal avalanches were characterized by the continuous activity patterns within the tissues. The calculating time bin ( $\Delta$ t) was set as 3 msec. Probability was calculated by following equation.

$$P(S) = kS^{-\alpha}$$

$P(S)$ is the probability of observing an avalanche of size $S$, $\alpha$ is the exponent of power law, giving the slope of relationship in a log-log coordinates, and $k$ is a proportionality.

##### *Optogenetic manipulation of the connected organoids*

To manipulate the activities of the connected organoids, we employed optogenetic tools (Fig. S2). AAV-CAG-hChR2$^{H134R}$-tdTomato was a gift from Karel Svoboda (Addgene plasmid # 28017). pAAV-CAG-ArchT-GFP was a gift from Edward Boyden (Addgene plasmid # 29777). Briefly, 5 µL of AAV virus prep were mixed with 500 µL of maintenance medium and then the mixture was replaced with the medium in the microfluidic device before at least 72h prior to the MEA measurement. Fiber-coupled LED of 470 nm (M470F3 - 470 nm, 17.2 mW (Min) Fiber-Coupled LED, 1000 mA, Thorlabs) for hChR2$^{H134R}$ and 565 nm LED (M565F3, 565 nm, 9.9 mW (Min) Fiber-Coupled LED, 700 mA, Thorlabs) for Arch-T were controlled by T-cube high power LED driver (LEDD1B, 1.2A, Thorlabs). Light was delivered to microfluidic device through multimode fiber (0.22 NA, High-OH, Ø105 µm Core, 250 - 1200 nm, Thorlabs). TTL pulse was generated by Arduino and it controlled LED driver. Source code was deployed and can be downloaded from https://github.com/TatsuyaOsaki/Arduino_optogenetics

***CRISPR-Cas9 Knock-in of GFP and mCherry fluorescent protein by electroporation***

To visualize axonal outgrowth, GFP or mCherry fluorescent protein were transfected to human PS cells. Each GFP and mCherry fluorescent protein were inserted to AAVS1 safe harbor locus. iPS cells were collected by TrypLE express treatment and centrifugation. Then, 5 μg of PX458-AAVS1 plasmid and 5 μg of AAVS1-Pur-CAG-EGFP plasmid (*21*) or 5 μg of AAVS1-Pur-CAG-mCherry plasmid (*21*) were mixed with $1\times10^6$ cells in 100 μL of Opti-MEM. The plasmid-cell mixture was transferred to NEPA cuvette (EC-002S) by a pipette and electrical pulses (Poring plus: voltage:125V, pulse length: 5 msec, pulse: 50 msec; a number of pulses: 2, decay rate 10%. Transfer pulse: voltage: 20V, pulse length: 50 msec, pulse: 50 msec; a number of pulses: 2, decay rate 40%,) were applied by a NEPA21 electroporator (NEPA gene). The electroporated iPS cells were seeded onto Matrigel-coated 4 wells of 6 well plate in mTeSR plus with 10 μM of Y-23632. After 24 h, transfected iPS cells were selected by adding 0.75 μg/ml of puromycin and treated for 2 days. Then, the cells were subcultured to Matrigel-coated 2 wells of 6 well plate and expanded. If necessary, iPS cells were further treated with puromycin or transfected iPS cells were manually selected by manual pipetting. PX458-AAVS1, AAVS1-Pur-CAG-EGFP, and AAVS1-Pur-CAG-mCherry plasmids were gifts from Drs. Adam Karpf and Su-Chun Zhang (Addgene 113194, 80945, and 80946).

***Cryosection and Immunocytochemistry***

Organoids were fixed in 4% paraformaldehyde (PFA) and 8% sucrose at 4°C for 15 min, washed 3 times with PBS (10 min incubation at RT for each wash), and transferred to 30% sucrose solution for incubation overnight at 4°C. Sucrose solution was then removed, and organoids were equilibrated with O.C.T compound at RT for 15 min. Organoids were then embedded within O.C.T compound on dry ice. Organoids were then stored at -80°C or cryosectioned to obtain 20 μm-thick slices. Cells were fixed with 4% paraformaldehyde for 20 min and then permeabilized with 0.2% Triton X-100 for 5 min. After blocking with 1% bovine serum albumin (BSA) for 2 h, the cells were incubated for 2 h at room temperature with a primary antibody. A secondary antibody was then administered for 2 h at room temperature. The primary antibodies were mouse anti-neuron-specific βIII tubulin (Biolegend 801202, 1:1200), rabbit anti-neuron-specific βIII tubulin (Sigma, ZooMAb, 1:200), mouse anti-human PAX6 (DHSB, 1:100), rabbit anti-human GAD67 (Santa Cruz, 1:100), rabbit anti-human vGluT1 (Sigma, ZooMAb, 1:200), mouse anti-human CTiP2 (Abcam, 1:100), rabbit anti-human SATB2(Abcam ab51502, 1:100), and rabbit anti-human MAP2 (Sigma, ZBR2290, 1:200). The secondary antibodies were Alexa Fluor 555 anti-rabbit IgG (H+L), Alexa Fluor 405 anti-rabbit IgG (H+L), Alexa Fluor 488 goat anti-mouse IgG (H+L), Alexa Fluor 488 goat anti-rabbit IgG (H+L), and Alexa Fluor 647 goat anti-rat IgG (H+L). Nuclei were stained with Hoechst dye for

20 min at room temperature, followed by three rinses with Dulbecco's Phosphate-Buffered Saline with $Ca^{2+}$ and $Mg^{2+}$ (D-$PBS^{++}$). All cells and samples were observed using a fluorescent microscope (Axio Observer, Zeiss) or a confocal laser scanning microscope (Zeiss).

##### Real-time reverse-transcription (RT)-PCR

To measure the biological activity of the cerebral organoids, total RNA was isolated from tissues with TriPure (Sigma) from the tissues. Reverse transcription was performed using a KOD One (Toyobo). The primer sequences are shown in **Table S1.** RT-PCR was performed with CFX Connect (BioRad) using KAPA SYBR FAST qPCR Master Mix (KAPA Biosystems). The mRNA level of glyceraldehyde 3-phosphate dehydrogenase (GAPDH) was used as the internal standard in all experiments. The RT-PCR experiments were repeated at least three times with cDNAs prepared from separate tissues.

##### *Simultaneous $Ca^{2+}$ imaging with customized microscopes*

To visualize neuronal activity by fluorescence, $Ca^{2+}$ indicator (GCaMP6f driven by CAG promoter) were transfected by using AAV1. pAAV.CAG.GCaMP6f.WPRE.SV40 was a gift from Douglas Kim & GENIE Project (Addgene #100836). Briefly, 5 µL of AAV1 virus prep were mixed with 500 µL of maintenance medium and then the mixture was replaced with the medium in the microfluidic device at least 3 days before measurement. AAV mixture was treated with 6-12 hours and then replaced with a fresh maintenance medium. On the day of the measurement, maintenance medium was replaced with Brainphys supplemented with 1% (v/v) B27 supplement with vitamin A, 1% (v/v) Glutamax, 20 ng/ml BDNF, and 1%(v/v) Penicillin/Streptomycin before 30 min measurement. Then, the device was set on the stage of customized microscopes. The configuration of the microscopes is described in Fig. S7. Time-lapse images were captured at a frame rate of more than 20 fps for 10 min. Data analysis was carried out using MATLAB (MathWorks). Regions of interest (ROIs) were manually drawn around the cell body of neurons in organoids. For each ROI time series, baseline fluorescence was defined as the average of the lowest 10% of samples. $\Delta F/F$ was computed as $(F - F0)/F0 \times 100$, where F is the instantaneous fluorescence from the raw ROI time series.

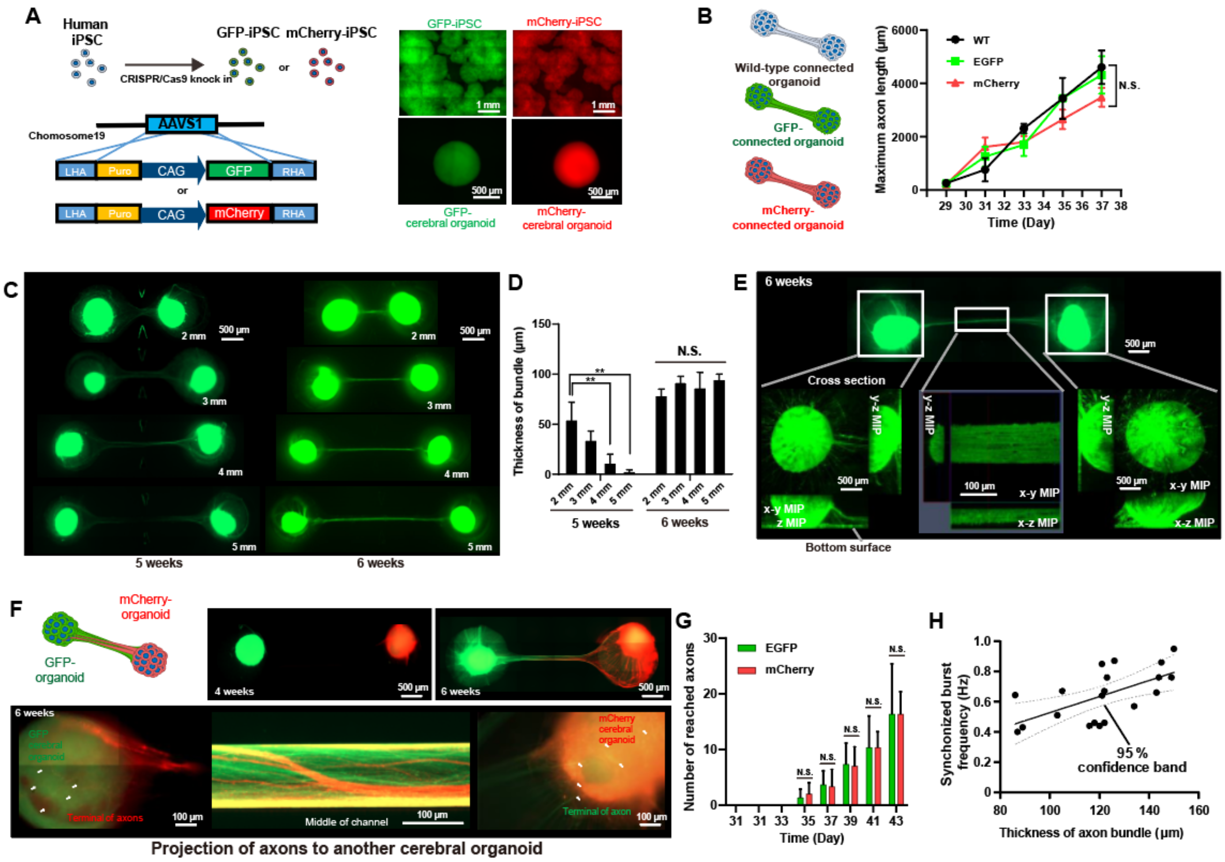


**Fig. S1 | Visualization of projected axons by CRISPR knock-in**

(A) CRISPR Knock-in of CAG promoter-driven EGFP and mCherry fluorescent tags into the AAVS1 safe harbor locus in human iPS cells. Then, GFP- or mCherry-labeled cerebral organoids were generated to visualize axon outgrowth in PDMS chips. (B) No significant phenotypic differences were observed between wild-type and EGFP- or mCherry-labeled cerebral organoids in terms of bundle formation speed. n = 3. (C) Tracking of axonal outgrowth of GFP-labeled connected cerebral organoids in different length of axon bundle (2, 3, 4, and 5 mm) at 5 and 6 weeks. (D) Thickness of axon bundles of GFP-labeled connected organoids at the center of micro channel. n = 3. (E) 3D confocal microscopic images of GFP-labeled connected organoid. (F) GFP-labeled and mCherry-labeled cerebral organoids were connected in a microfluidic device. GFP- and mCherry labeled axons projected to another mCherry and GFP-labeled cerebral organoid via merged thick axon bundle after two weeks in the chip. (G) Number of axons that reached the other cerebral organoid in a chip. n = 3. (H) Correlation between thickness of axon bundle and synchronized burst frequency. Burst frequency increased with increasing the axon bundle thickness. **, $p<0.01$, one-way ANOVA or student's t-test. Error bars, ±SD.

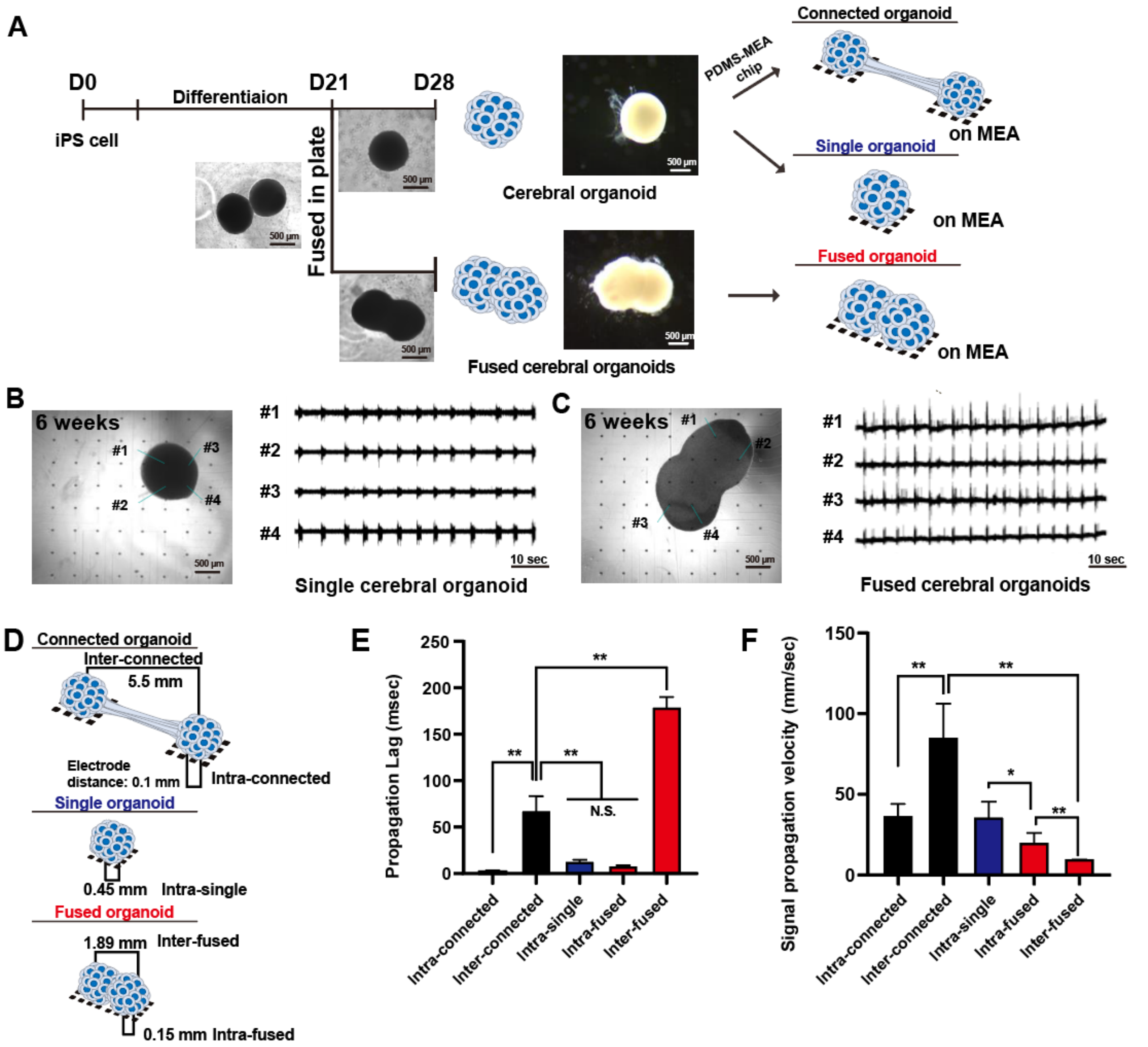


**Fig. S2 | Comparison of single, fused, and connected organoids.**
(A) Schematic illustration of generating single, and fused and connected organoids. All cerebral organoids were generated in the same manner up to 21 days. To generate fused organoids, two cerebral organoids were placed into a well of low-adherent 96 well plate. Single and fused organoids were then placed onto MEA probes on day 28. Organoids were introduced into a PDMS-MEA chip for generating connected organoid on day 28. After 2 weeks of culture on MEA probes, neuronal activities were measured. (B) A representative image of single organoid on a MEA probe. Periodic and synchronized neuronal activity was observed. (C) A representative image of fused organoids on a MEA probe. More aggressive periodic and synchronized neuronal activity than single organoid was observed. (D) Illustration of intra- and inter -organoid activity lag detection by electrodes. (E) Quantification of propagation lag between electrodes. n = 4. 'Inter-connected' exhibited significantly smaller lag than 'inter-fused'. (F) Signal propagation velocity. 'inter-connected' signal transduction was faster than others. n = 4 *, p<0.05, **, p<0.01, one-way ANOVA or student's t-test. Error bars, ±SD.

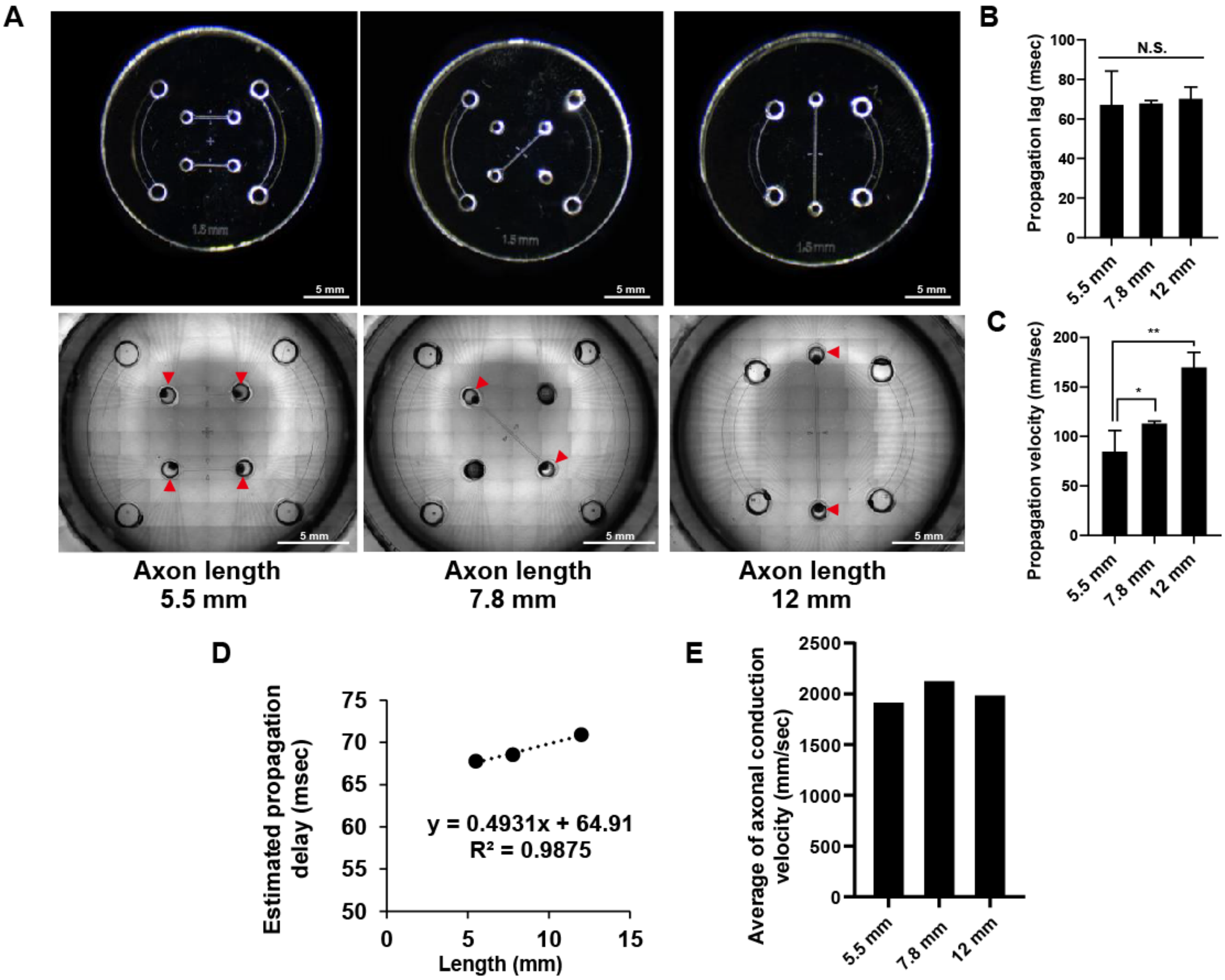


**Fig. S3 | Organoids connected with various length of axon bundles.**
(A) Three types of microfluidic chips with different lengths of channels (5.5 mm. 7.8 mm, and 12 mm). Red arrowheads indicate organoids placed in the chips. (B) Propagation lag between the connected organoids. n = 4. (C) Propagation velocity in different length of axon bundle. n = 4. (D) To estimate axon conduction velocity, the propagation delay was plotted over distance (mm). Delay constant was 65 msec. (E) Actual axonal conduction velocity were equally estimated as around 2000 mm/sec. *, $p<0.05$, **, $p<0.01$, by one-way ANOVA or student's t-test. Error bars, ±SD

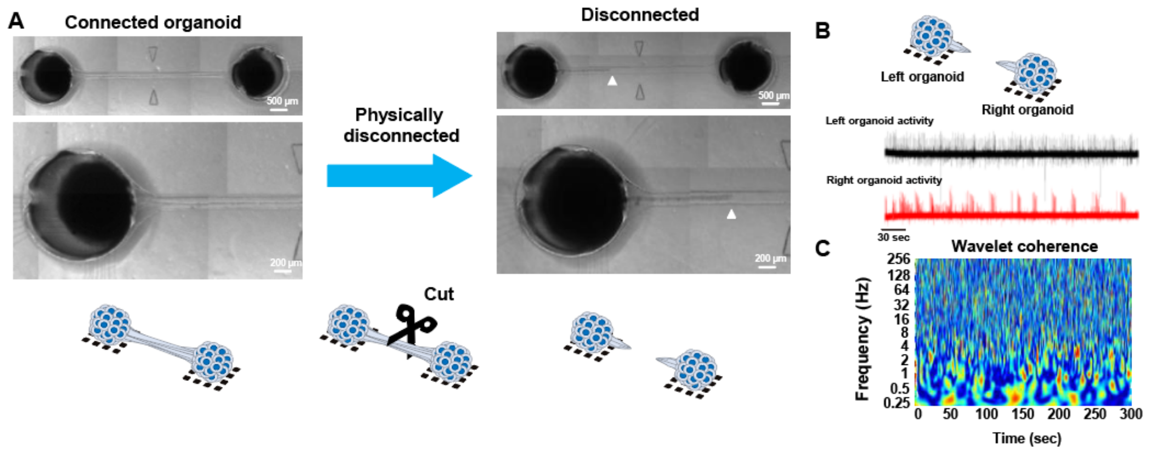


**Fig. S4 | Physically disconnected organoids lose synchronized activity.**
(A) An axon bundle between connected organoids was physically cut and disconnected (white arrow). (B) After cutting the axon bundle, synchronized activity was completely lost. (C) Wavelet coherence indicated little or no synchronized activity in the disconnected organoid.

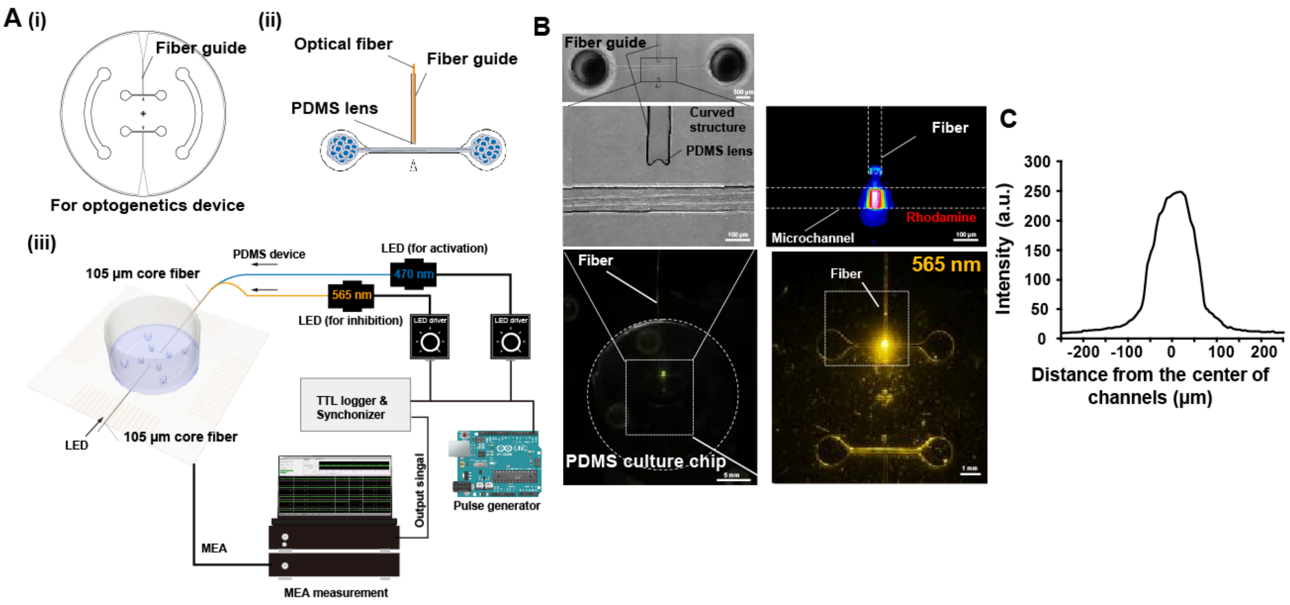


**Fig. S5 | Optogenetics set-up for activation and inhibition of neuronal activity in connected organoids.**

(A) (i) A microfluidic device for optogenetic control. A fiber guide (thin channel) positioned an optical fiber. (ii) An optical fiber was positioned perpendicular to the axon bundle with a 100 µm gap. (iii) An optical fiber was connected to a 470 nm or 565 nm LED and a pulse generator (Arduino). MEA measurement was conducted. Light exposure timing and a representative channel from MEA amplifier were recorded in TTL logger to synchronize TTL signal and recoding. (B) An axon bundle and an optical fiber. Curved structure serves as a PDMS lens which helps the light to be focused on the axon bundle (C).

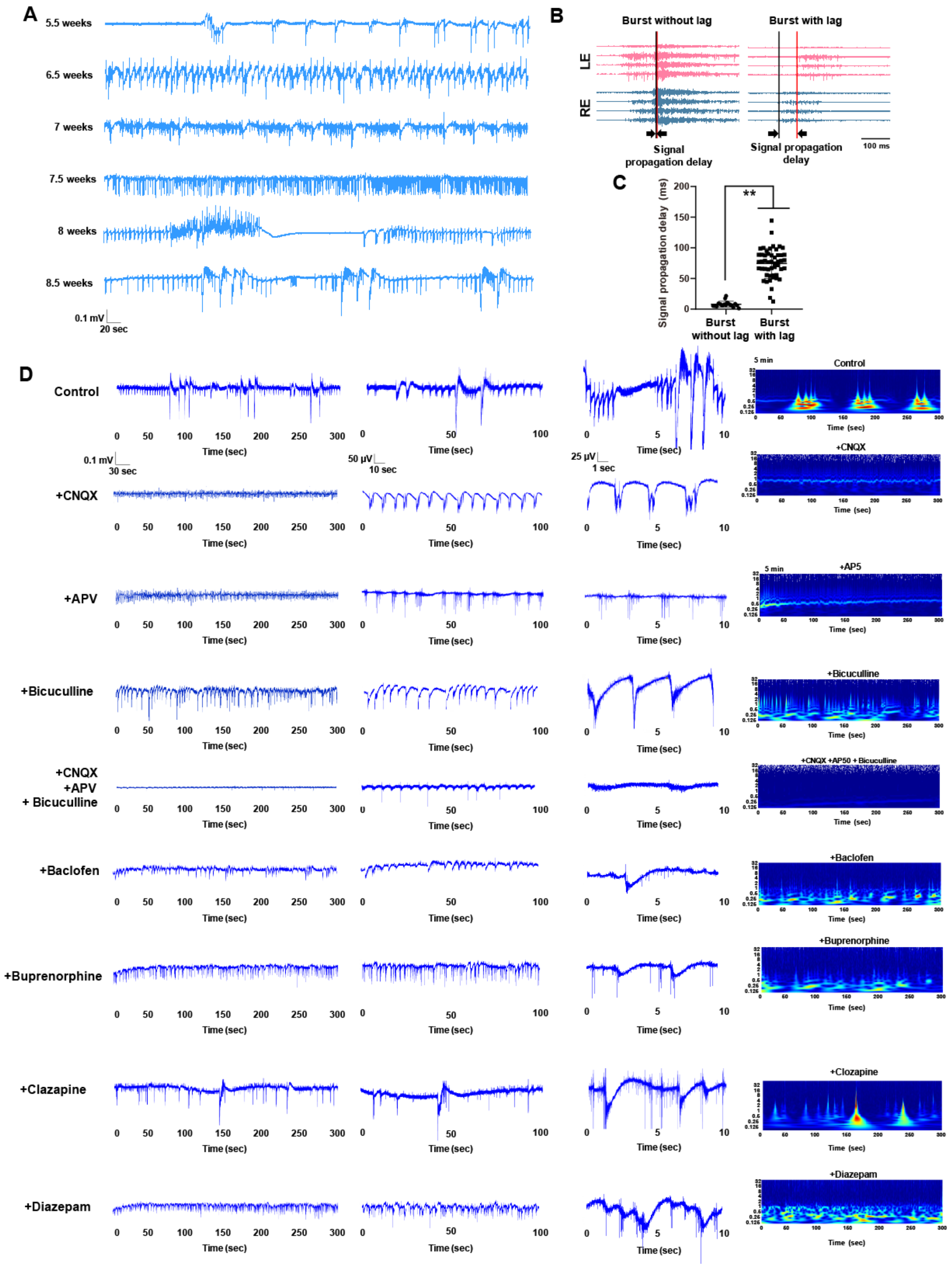


**Fig. S6 | Connected organoid maturation and drug treatment**

(A) LFP signals of the connected organoids in different culture periods. (B, C) Burst events are observed with and without time lag between the connected organoids in 9 weeks. (D) LFP signal and scalorgram of wavelet transformation with various drugs.

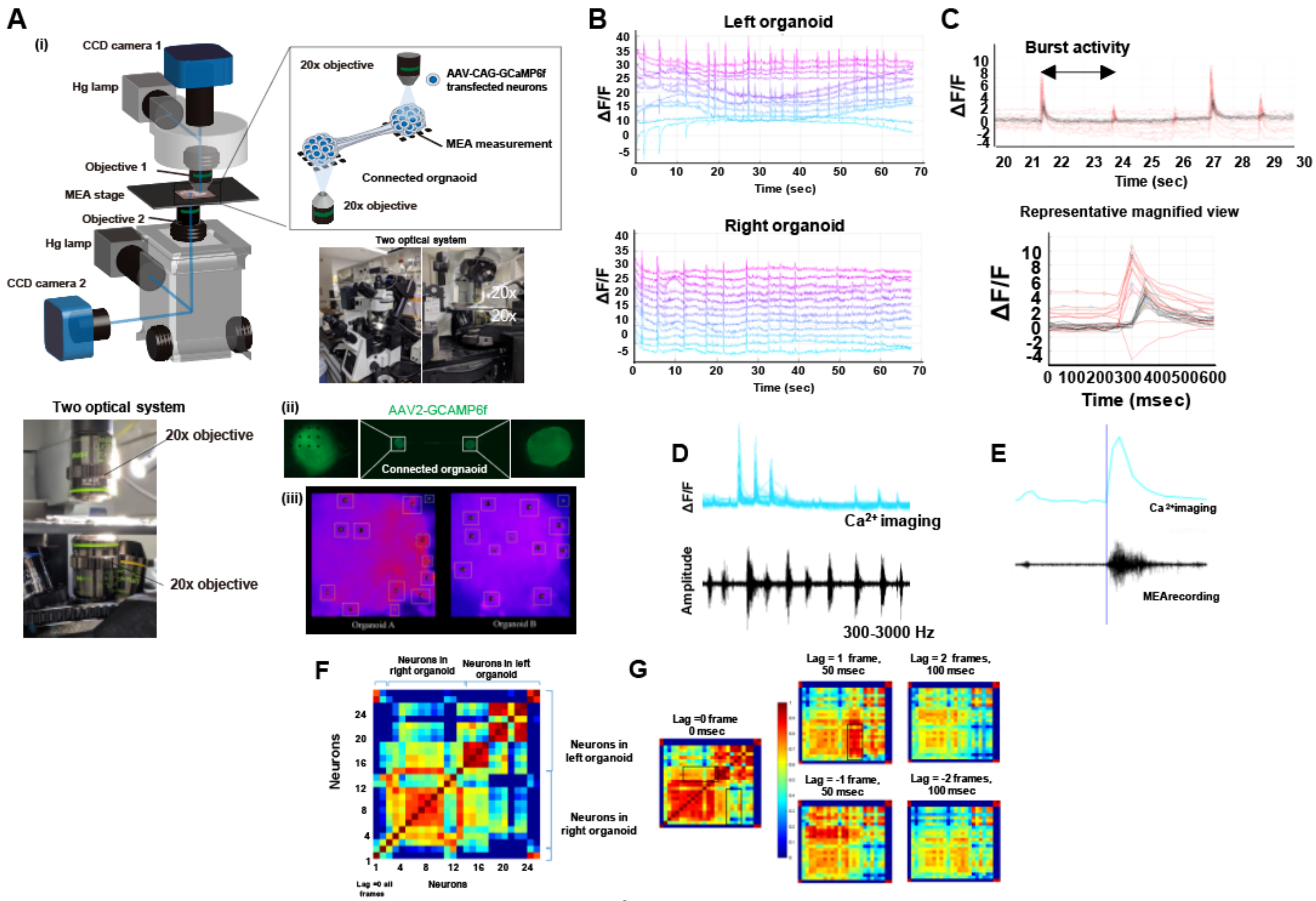


**Fig. S7 | Simultaneous measurement of $Ca^{2+}$ transient and electrical activity**

(A) (i) Optical set-up for simultaneous $Ca^{2+}$ imaging and MEA recording of the connected organoids. (ii, iii) $Ca^{2+}$-reporter gene (GCaMP6f) was transiently transfected to the connected organoids 3 days prior to the measurement by AAV2. Then, time-series images were captured with a custom-made microscope setup while LFP activity form MEA was acquired simultaneously. (B) The firing patterns of the cells in the connected organoids shown by a trace image of the calcium response at 7 weeks of differentiation. (C) Synchronized burst activity between left (red line) and right (black line) organoids was observed. Burst delay between the two connected organoids (~50 msec) was consistent with MEA recording result. (D) Plot of both $Ca^{2+}$ activity and MEA signal. Two type of signals corresponded each other. (E) Consistency of $Ca^{2+}$ imaging and MEA recording during a burst activity. (F, G) Correlation matrix from 12 neurons on left organoid and 12 neurons on right organoids is shown for 10 sec measuring time. 2 negative control ROIs serve as the reference. $Ca^{2+}$ activity was strongly correlated within each organoid. $Ca^{2+}$ activity was more strongly correlated between the organoids when the signals were shifted 50 msec (1 frame).

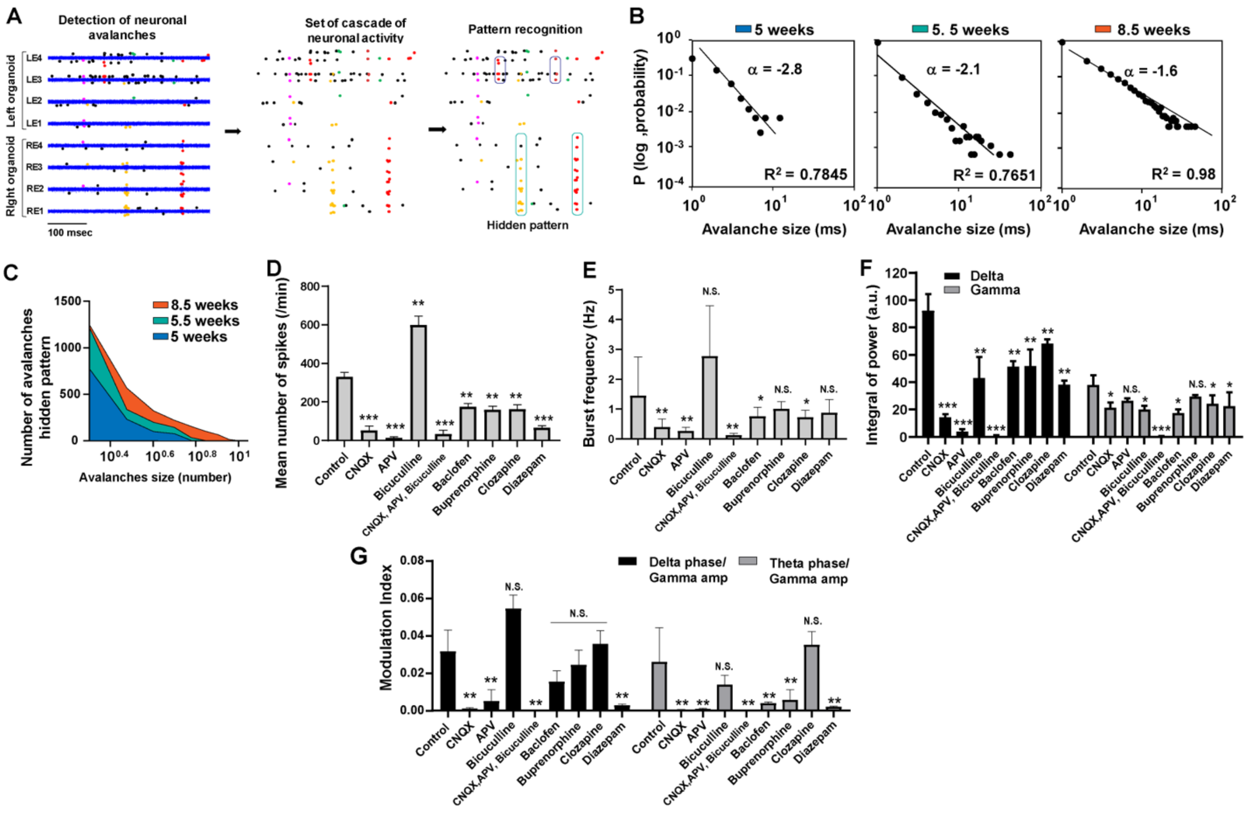


**Fig. S8 | neuronal avalanches and neuromodulator compounds reveal complex network activity in the connected organoids**

(A) Schematic illustration of analyses of neuronal avalanches including the extraction of neuronal avalanche cascades and hidden pattern recognition by Markov model. Neuronal avalanches were calculated from 8 electrodes. The cascade of single spikes was characterized at 3 msec scale size. (B The log plot of neuronal avalanche size and probability at 5, 5.5, and 8.5 weeks of culture. (C) The number of hidden patterns in neuronal avalanches at 5, 5.5, and 8.5 weeks. (D, E, F, G) Comparison of activity patterns of the connected organoids upon treatment with neuromodulator compounds. Mean number of spikes (D), burst frequency (E), power integral in delta band (F), and PAC modulation index (G) were assessed. *p<0.05, **p<0.01; one-way ANOVA. Error bars indicate SD.

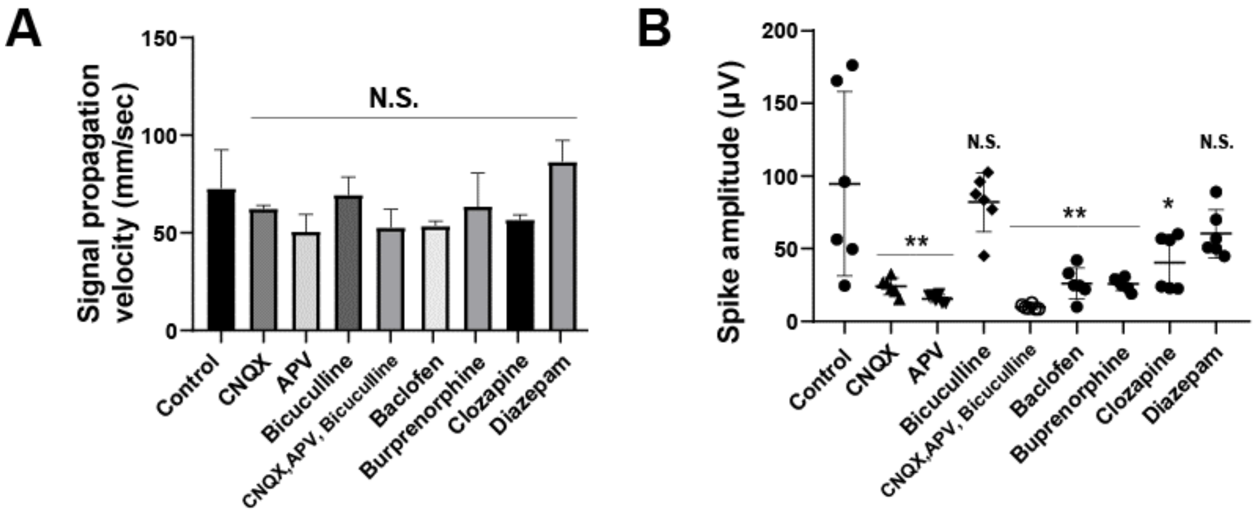


**Fig. S9 | Signal propagation velocity and spike amplitude**

(A) Signal propagation velocity in the presence of drugs. Drugs did not affect the velocity. n = 4. (B) Spike amplitude in the presence of drugs. n = 4. *, $p<0.05$, **, $p<0.01$, one-way ANOVA or student's t-test. Error bars, ±SD.

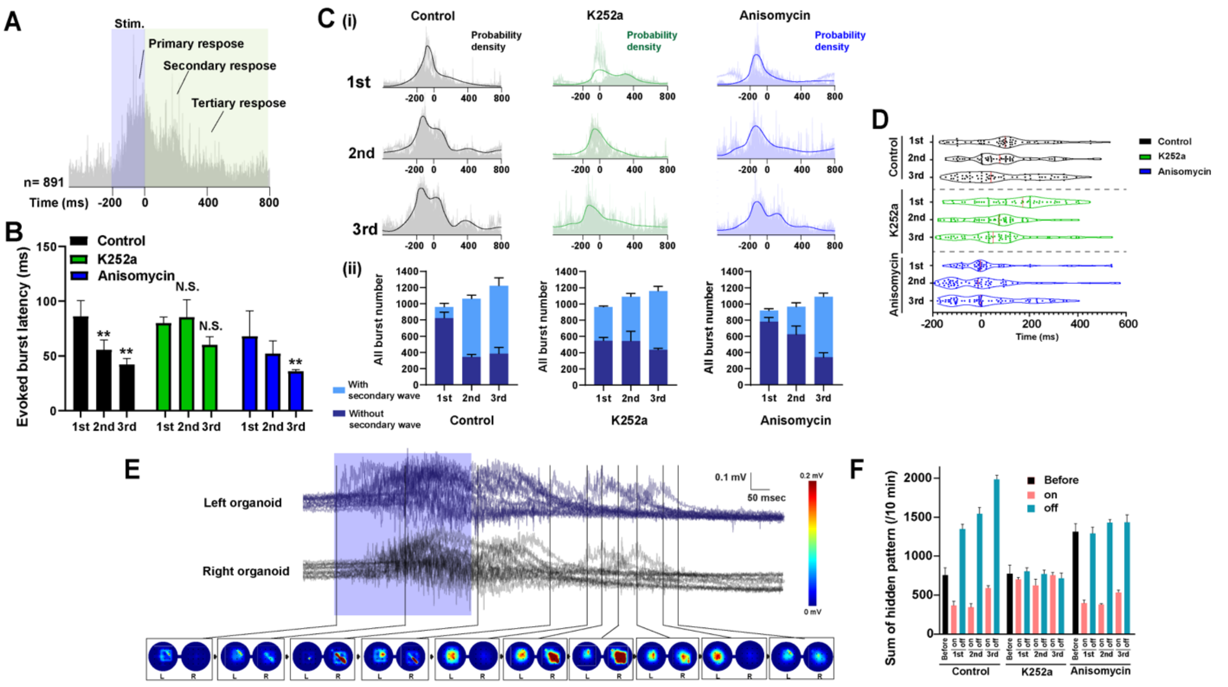


**Fig. S10 | Diverse evoked burst patterns supported by CaMKII-dependent signaling**

(A) Representative image of sorted evoked bursts by optogenetic stimulation and color-coded map depicting the voltage distribution from neurons in left and right cerebral organoids. A total of 891 burst traces is presented. Light stimulation induced evoked spikes, and the burst waves persisted after light stimulation. Secondary, and tertiary waves were also observed. (B) Latency of the evoked bursts. Repeated light stimulation (at second and third attempts) significantly decreased evoked burst latency in control and anisomycin-treated conditions, whereas K252a treatment did not influence the latency of evoked bursts. n = 3. (C) (i) Overlayed power histograms of evoked bursts and kernel density estimation (line) in the presence of K252a and anisomycin. Repeated light stimulation increased burst wave complexity. (ii) Percentage of evoked bursts with and without secondary peaks. (D)Violin plots of burst wave peak times. n = 50. (E) Representative crosstalk between the two connected organoids in self-evoked bursts. (F) Less hidden patterns were observed during the light-on than during the light-off period. K252a treatment suppressed the number of hidden patterns even after repeated training. *p<0.05, **p<0.01; one-way ANOVA or student’s t-test. Error bars indicate SD.

**Table S1. Real-time PCR primers**

| Gene | Forward primer 5'-3' | Reverse primer 5'-3' |
|---|---|---|
| **DCX** | TGCCAGAAAGTCTCAACAGCC | GCGTACACAATCCCCTTGAAGTA |
| **FOXG1** | ATGATGCAAGAATCTGGGACTG | AGGAGGGCGAGAAGAAGAAC |
| **GAD1** | TGGCGTTTCTGCAAGATGTTA | CACAAGGCGACTCTTCTCTTC |
| **GAD2** | GGCCGCAACCAAAATTCAGG | TTGGTCTGCCAATTCCCAATTAT |
| **Gria1** | CGA AAA TGC CAG CTG ATA TAA | CTT CCC GGA CCA AAG TGA TAG |
| **LHX6** | CAG GAG ACA GAG ATG AGA ATT CC | TGAACGGGGTGTAGTGGATGT |
| **MAP2** | GTC TCA GGA CAG TGC TGA GCC TTC | CAG GAG TGA TGG CAG TAG AC |
| **Nestin** | GAC GCA ATG GTT CAG CCT TTT | TCC CCT GAG GAC CAG GAG TCT C |
| **NKX2.1** | GCCGTACCAGGACACCATG | GTGTCACGAGAAGTAGAGGTCT |
| **PAX6** | ACCCATTATCCAGATGTGTTTGCCCGAG | ACAGGCATCTGAGGTGAACAG |
| **SATB1** | GATCATTTGAACGAGGCAACTCA | TGGACCCTTCGGATCACTCA |
| **SATB2** | GACAGTGGCCGACATGCTAC | ATGTTCAAAGAACTCGTGGCA |
| **SOX2** | GCCGAGTGGAAACTTTTGTCG | CAAAGTCCTGATACCAGCATCTT |
| **TBR1** | CACCGCCACCAAACTGAGAT | TCCCGAGAGAGGAATTAGAACTT |
| **TBR2** | GGCCAAGGGTCACTACACG | CGAACACATTGTAGTGGGCAG |
| **TUJ1** | TCA ATA ACA GCA CGA CCC AC | GCAGTCGCAGTTTTCACACTC |
| **vGluT1** | TGG TCG TTG GCT ATT CTC ATA C | TCC TGG AAT CTG AGT GAC AAT G |
| **GAPDH** | TGT GGG CAT CAA TGG ATT TGG | ACA CCA TGT ATT CCG GGT CAA T |

**Movie S1.**

Simultaneous detection of $Ca^{2+}$ transient from two interconnected organoids.
